# SCReadCounts: Estimation of cell-level SNVs from scRNA-seq data

**DOI:** 10.1101/2020.11.23.394569

**Authors:** NM Prashant, Nawaf Alomran, Yu Chen, Hongyu Liu, Pavlos Bousounis, Mercedeh Movassagh, Nathan Edwards, Anelia Horvath

## Abstract

Recent studies have demonstrated the utility of scRNA-seq SNVs to distinguish tumor from normal cells, characterize intra-tumoral heterogeneity, and define mutation-associated expression signatures. In addition to cancer studies, SNVs from single cells have been useful in studies of transcriptional burst kinetics, allelic expression, chromosome X inactivation, ploidy estimations, and haplotype inference. To aid these types of studies, we have developed a tool, SCReadCounts, for cell-level tabulation of the sequencing read counts bearing SNV reference and variant alleles from barcoded scRNA-seq alignments. Provided genomic loci and expected alleles, SCReadCounts generates cell-SNV matrices with the absolute variant- and reference-harboring read counts, as well as cell-SNV matrices of expressed Variant Allele Fraction (VAF_RNA_) suitable for a variety of downstream applications. We demonstrate three different SCReadCounts applications on 59,884 cells from seven neuroblastoma samples: (1) estimation of cell-level expression of known somatic mutations and RNA-editing sites, (2) estimation of celllevel allele expression of germline heterozygous SNVs, and (3) a discovery mode assessment of the reference and each of the three alternative nucleotides at genomic positions of interest that does not require prior SNV information. For the later, we applied SCReadCounts on the coding regions of *KRAS*, where it identified known and novel recurrent somatic mutations in a low-to-moderate proportion of cells. The SCReadCounts read counts module is benchmarked against the analogous modules of GATK and Samtools. SCReadCounts is freely available (https://github.com/HorvathLab/NGS) as 64-bit self-contained binary distributions for Linux and MacOS, in addition to Python source.

## Introduction

Single cell RNA sequencing (scRNA-seq) brings major advantages over bulk RNA-seq analyses, especially the ability to distinguish cell populations and to assess cell-type specific phenotypes [1–3]. Connecting these phenotypes to cell-level genetic variants (such as Single Nucleotide Variants, SNVs) is essential for phenotype interpretation. In cancer, studies on cellular genetic heterogeneity have been instrumental in tracing lineages and resolving subclonal tumor architecture [3–18]. In addition to cancer studies, SNV observations from single cells have been useful in studies of transcriptional burst kinetics, allelic expression, chromosome X inactivation, ploidy estimations, haplotype inference, and quantitative trait loci (QTL) [4–6,11,14–17,19–21].

To aid with the considerable data-analysis and data-management demands of such studies, we have developed a tool, SCReadCounts, for cell-level quantitation of SNV expression. Provided with barcoded scRNA-seq alignments and genomic loci and alleles of interest, SCReadCounts tabulates, for each cell, the reference and variant read counts (n_ref_ and n_var_, respectively), and expressed Variant Allele Fraction (VAF_RNA_ = n_var_ / (n_var_ + n_ref_)). SCReadCounts generates a cell-SNV matrix with the absolute n_var_ and n_ref_ counts, and a cell-SNV matrix with the VAF_RNA_ estimated at a user-defined threshold of minimum number of required sequencing reads (minR) (Figure 1a and S_Fig.1). Particular strengths of SCReadCounts include its named, explicit, flexible, and configurable read-filtering and cell-barcode extraction capabilities; accounting for all reads overlapping each locus, whether counted or ignored; and straightforward input and output formats – these features make SCReadCounts easy to integrate in multi-tool data-analysis pipelines. The cell-SNV matrices then can be used for a wide range of downstream analyses.

**Figure 1.**
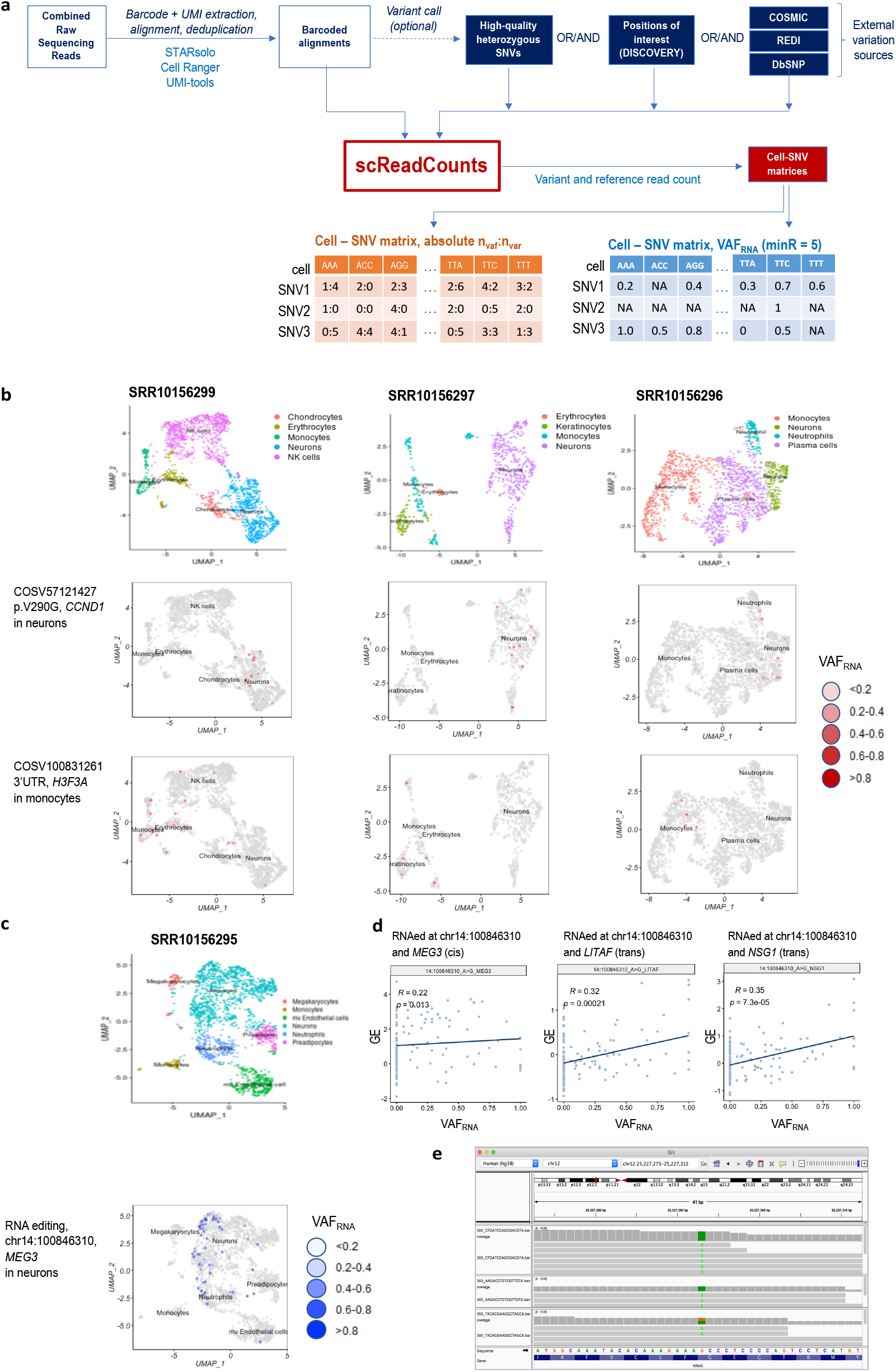
**a.** SCReadCounts workflow using publicly available tools. **b (top)** Twodimensional UMAP clusters showing different cell types from samples SRR10156299, SRR10156297 and SRR10156295; **(middle)** Quantitative visualization (red) of the VAF_RNA_ of somatic mutations in the corresponding tumor samples: COSV57121427 in *CCND1* is primarily expressed in neurons, while COSV100831261 in *HSF3A* **(bottom)** is primarily expressed in non-neuronal cells, including monocytes. **c.** Two-dimensional UMAP clusters showing cells classified by type **(top)** and visualizing RNA-editing levels (blue) in the gene *MEG3*, where the intensity of the color corresponds to the proportion of edited reads (bottom, sample SRR10156295). **d (left)** Cis-scReQTL correlation between edited levels (x-axis) and GE (y-axis) of *MEG3.* **(middle and right):** trans-scReQTL between editing levels (x-axis) in *MEG3* and GE of *LITAF* and *NSG1* GE (y-axis); the sample is SRR10156295. **e**. IGV visualization of variable VAF_RNA_ of the novel somatic mutation Gly77Gly (12:25227293_G>A) in the gene *KRAS* in three individual cells of sample SRR10156300.

Unlike variant callers (i.e. GATK, Samtools [22,23]), SCReadCounts estimates the read counts per allele and per cell, across all cells, including cells in which the position of interest is covered with only reference reads. This is particularly useful in scRNA-seq settings, where it enables distinguishing cells with monoallelic reference expression from those with no gene expression. The later can be used to assess cell-level allele dynamics, and to correlate variant expression to gene expression [12].

## Results

We have explored a variety of SCReadCounts applications on over 300,000 single cells from 6 different studies on normal and tumor human samples, including adipose tissue, adrenal neuroblastoma, acute myeloid leukemia, non-small lung cancer, prostate carcinoma, and the MCF7 cell line derived from breast adenocarcinoma [12,24–30]. Here we demonstrate three different SCReadCounts applications on 59,884 cells derived from seven neuroblastoma samples [26]: (1) estimation of cell level expression of known somatic mutations and RNA-editing sites, (2) estimation of cell level allele expression from biallelic positions as called in the pooled scRNA-seq data, and (3) a discovery mode assessment of the reference and each of the three alternative nucleotides at genomic positions of interest. The discovery mode does not require prior knowledge on existing genetic variants and is particularly convenient for a quick focused assessment of a gene or a group of genes or regions of interest.

For all three applications the scRNA-seq data was processed using publicly available tools. In the exemplified workflow (Figure 1a), the raw sequencing reads are aligned to the reference human genome (GRCh38) using STARsolo (v.2.5.7b) which processes the cellular barcodes and deduplicates the alignment retaining the reads with the highest mapping quality using the unique molecular identifiers (UMI) [31]. In addition to STARsolo, SCReadCounts accepts barcoded alignments generated by CellRanger and UMI-tools [32,33]. The alignments can be filtered to correct for allele-mapping bias by removing reads mapped ambiguously due to the variant nucleotide (WASP); this filtering utilizes the same list of positions to be used as input for SCReadCounts [34].

### SCReadCounts on known variant loci

SCReadCounts can be applied to assess known genetic variation loci such as somatic mutational hotspots or RNA-editing sites. We first asked if we could assess known somatic mutations in a set of seven neuroblastoma scRNA-seq samples (S_Table 1) [26]. Known somatic mutations (118,394 loci) were extracted from COSMIC by selecting loci in Tier 1 Cancer Census Genes not present in dbSNP, in order to ensure germline variants are excluded (dbSNPv.154). These somatic mutations (S_Table 2) were provided to SCReadCounts. A minimum of 3 sequencing reads (n_var_ + n_ref_ >= 3, minR = 3) was required for loci to be considered for further analysis. SCReadCounts identified 443 distinct COSMIC mutations expressed in at least 4 individual cells in one or more of the 7 neuroblastoma samples (S_Table 3). Examples include COSV57121427 in the coding region of *CCND1* and COSV100831261 in the 3’UTR of *H3F3A* (Figure 1b). Notably, COSV57121427 was almost exclusively expressed in neurons, while COSV100831261 was preferentially expressed in non-neuronal cells of the same samples; these observations were consistent across neuroblastoma tumors from different patients.

Next, we demonstrate that SCReadCounts can quantify cell-specific RNA-editing in the same neuroblastoma samples. We use the previously described single nucleotide RNA-editing events catalogued in the REDI database [35], after excluding genomic positions in repetitive regions or that coincide with a potential germline variant. A total of 107,053 distinct RNA-editing sites were provided to SCReadCounts (S_Table 4) along with the corresponding scRNA-seq alignments. At minR = 5, SCReadCounts identified 72 positions which were edited in at least 2 cells in one or more of the 7 neuroblastoma samples (S_Table 5). We investigated the A>G RNA-editing event at 14:100846310 in the cancer-implicated lincRNA *MEG3;* this position was edited in 6 of the 7 samples. First, we observed that cells with *MEG3* RNA-editing were seen predominantly in neurons, where they displayed clear positional clustering (Figure 1c). Second, the proportion of edited RNA molecules (as reflected through the VAF_RNA_), suggest variable degrees of RNA editing per cell. We then checked whether the degree of RNA-editing was correlated with gene expression using scReQTL [12]. This analysis showed weak positive correlation of the degree of editing at 14:100846310 and the expression of the harboring *MEG3* (cis-scReQTL, Figure 1d left), as well as stronger correlation with the expression of other genes (trans-scReQTLs), including the cancer-implicated *LITAF* and *NSG1* (Figure 1d middle and right).

### SCReadCounts after variant call

SCReadCounts can be applied in conjunction with variant callers to estimate the cell-specific allele expression of germline heterozygous SNVs. To explore this application, we called variants from the pooled alignments using GATK (v4.1.7.0, [22]), and filtered the calls retaining high quality biallelic positions for which both the variant and the reference allele were supported by a minimum of 50 sequencing reads, as we have previously described [12]. The variant lists were then provided to SCReadCounts together with the corresponding alignments. The resulting VAF_RNA_ estimates can be used to explore the cell-level allelic expression. For example, the distribution of the VAF_RNA_ at minR = 5 across the cells for each of the seven neuroblastoma samples is plotted on S_Figure 1, which shows that many of the SNVs have skewed or mono-allelic expression. VAF_RNA_ estimates for positions covered by at least 10 total reads (minR = 10) in 20 and more cells per sample are summarized in S_ Table 6. A systematic analysis of the distribution VAF_RNA_ of heterozygous SNVs at different minR is analysed in our recent work [30]. In addition, VAF_RNA_ estimates can be used to explore correlations between allele- and gene-expression in single cells using scReQTL [12]. Our previous research applying scReQTL on normal adipose datasets has shown that scReQTLs are substantially enriched in GWAS-significant SNVs and in known gene-gene interactions [12].

### SCReadCounts in discovery mode

As mentioned earlier, SCReadCounts can be applied in a discovery mode which does not use any prior knowledge of SNVs. In this use-case, SCReadCounts considers positions of interest where the reference nucleotide is substituted with each of the three alternative nucleotides. Such SCReadCounts inputs can be generated for a gene, region or a group of genes/regions of interest, either manually, or using a script supplied with SCReadCounts. Herein, we demonstrate this approach using an enumeration of each position in the coding region of *KRAS* (S_Table 7), mutations in which have been implicated in neuroblastoma. Across the seven samples, SCReadCounts identified a total of 30 distinct mutations that do not coincide with known germline variants in the coding sequence of *KRAS* (S_Table_8). The mutations included missense, nonsense and synonymous substitutions in up to 15 individual cells per sample. Eight of the 30 mutations were seen in more than one sample; for example, the synonymous substation 12:25227293_G>A (Gly77Gly, Figure 1e) was seen in 4 out of the 7 samples. Seven of the 30 mutations were previously catalogued in the COSMIC database – the remaining 23 substitutions represent novel *KRAS* variants.

### Performance

We compared the variant and reference read counts tabulations of SCReadCounts with the analogous modules of the mpileup utility of Samtools [23] and the haplotype caller of GATK. SCReadCounts default options generate nearly identical values to mpileup and GATK (Figure 2). SCReadCounts uses, by default, a very simple read-filtering criteria, but it can also be readily configured to achieve scenario-specific, mpileup-consistent, or GATK-consistent results, with optional explicit output of the number of reads discarded by each filtering rule. In regard to efficiency, on our system (2×14 cores CPUs with 1.5TB RAM compute node) processing of a file containing ~5000 cells, ~150mln reads, and ~80K SNVs, requires approximately 4h for the tabulation of n_var_ and n_ref_, and up to 2 minutes for the generation of the cell-SNV matrices. The later enables the users to quickly generate VAF_RNA_ matrices at various minR.

**Figure 2.**
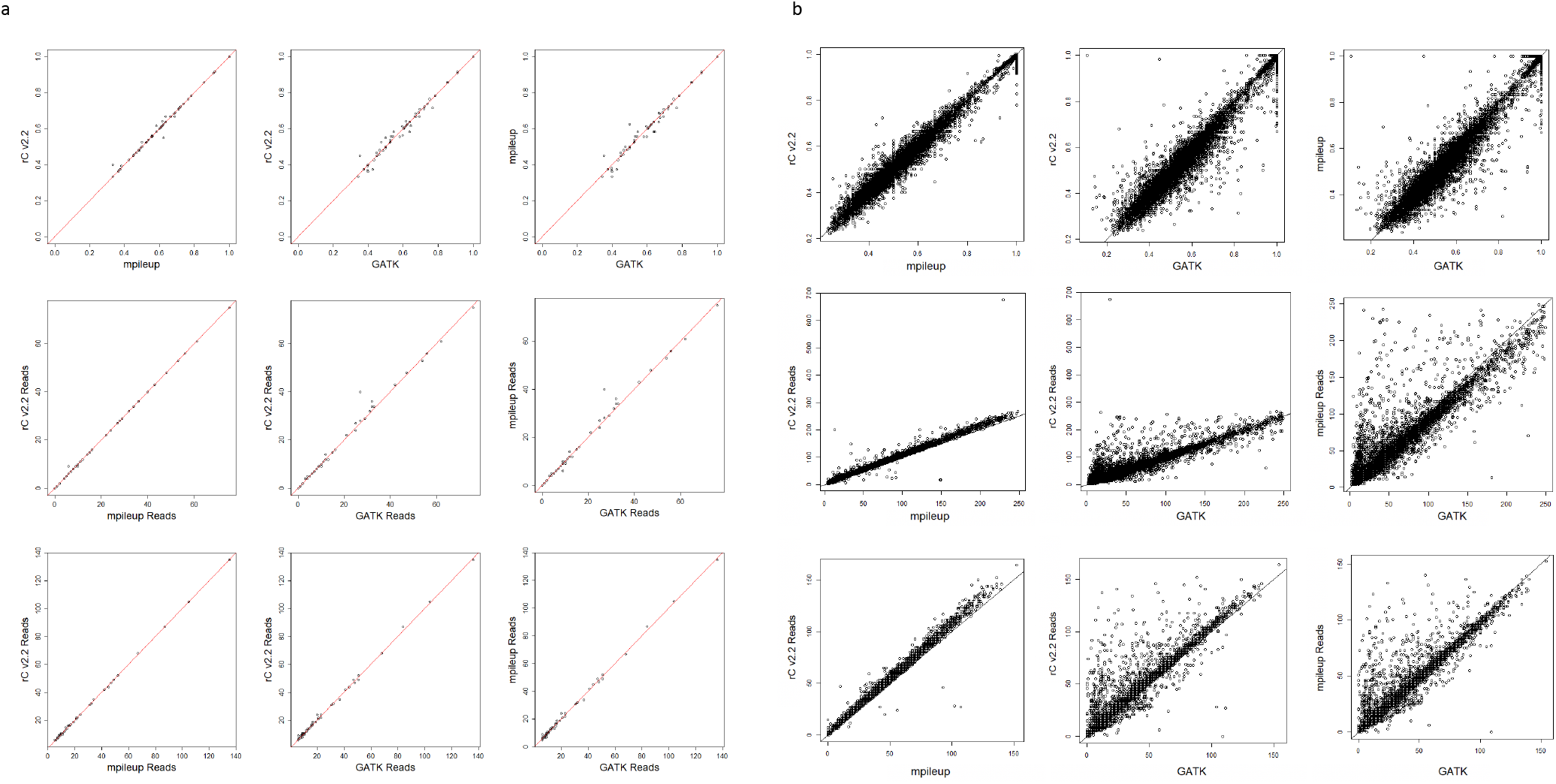
Concordance between n_var_, n_ref_ and VAF_RNA_ estimates across SCReadCounts (rc_v.2.2), the mpileup utility of Samtools, and the haplotype caller of GATK in individual cell scRNA-seq alignments (a) and pooled scRNA-seq alignments (b) of sample SRR10156295. Top: VAF_RNA_ middle: n_var_, bottom: n_ref_.

## Discussion

SCReadCounts supports several use-cases specific to scRNA-seq. First, and perhaps most importantly, SCReadCounts can detect and quantify SNVs present in a low proportion of cells, such as somatic and mosaic SNVs. SNVs present in a low proportion of cells are known to be frequently missed by variant calls carried out on pooled scRNA-seq alignments [36,37]. Indeed, the vast majority of the somatic mutations detected by SCReadCounts were not called on the pooled scRNA-seq data by GATK at its default setting (See S_Tables 3 and 8). In the herein presented analysis, SCReadCounts detected novel somatic mutations occurring in individual cells or in a small number of cells in one or more of the neuroblastoma samples. Second, SCReadCounts provides per cell quantitation of absolute and relative number of variant reads across all cells, including those where the position is covered with only reference reads, allowing the identification of cells with preferential expression of the variant or the reference allele. Furthermore, the flexible VAF_RNA_ minR enables tuning of SCReadCounts to the particular application and depth of sequencing. Here, a major consideration is the balance between inclusivity (low minR) and higher-confidence VAF_RNA_ estimates (high minR). In our analyses we use different minR depending on the application [12,30]. For example, to confidently estimate allele expression of germline variants in highly transcribed genes, high minR is needed. In contrast, assessments of somatic mutations would benefit of high inclusivity using low minR. Finally, SCReadCounts generates cell-SNV matrices that are analogous to the cell-gene matrices generated by popular scRNA-seq tools, which streamlines downstream applications combining SNV and gene expression. We have explored correlating SNVs to gene expression for heterozygote germline variants at single cell level using scReQTL [12]. Here, we apply scReQTL on RNA-editing sites and identify cis- and trans-correlations between RNA-editing and gene expression (see Figure 1d). In conclusion, we believe that SCReadCounts supplies a fast and efficient solution for estimation of scRNA-seq genetic variance.

## Ethics approval and consent to participate

The study uses only previously published and freely available datasets.

## Availability of data and materials

All the data analyzed in this study are supplied with the supplemental material or available as indicated in the cited publications.

## Authors’ contributions

PNM, NA, YC and NE developed and tested the software; HL, PB and MM performed the analyses and contributed to the visualization; NE conceptually developed and implemented the tool, AH devised and supervised the study and wrote the manuscript.

